# Hybridization and postzygotic isolation promote reinforcement of male mating preferences in a diverse group of fishes with traditional sex roles

**DOI:** 10.1101/325498

**Authors:** Rachel L. Moran, Muchu Zhou, Julian M. Catchen, Rebecca C. Fuller

## Abstract

Behavioral isolation is thought to arise early in speciation due to differential sexual and/or natural selection favoring different preferences and traits in different lineages. Alternatively, behavioral isolation can arise due to reinforcement favoring traits and preferences that prevent maladaptive hybridization. In darters, female preference for male coloration has been hypothesized to drive speciation, because behavioral isolation evolves before F1 inviability. However, as with many long-lived organisms, the fitness of second generation hybrids has not been assessed because raising animals to adulthood in the lab is challenging. Recently, reinforcement of male preferences has been implicated in darters because male preference for conspecific females is high in sympatry but absent in allopatry in multiple species pairs. The hypothesis that reinforcement accounts for behavioral isolation in sympatry assumes that hybridization and postzygotic isolation are present. Here, we used genomic and morphological data to demonstrate that hybridization is ongoing between orangethroat and rainbow darters and used hybrids collected from nature to measure postzygotic barriers across two hybrid generations. We observed sex ratio distortion in adult F1s and a dramatic reduction in backcross survival. Our findings indicate that selection to avoid hybridization promotes the evolution of male-driven behavioral isolation via reinforcement in this system.

## Introduction

The increasing availability of genomic sequence data for non-model organisms has revealed that hybridization is surprisingly common between species (Mallet 2005; Abbott et al. 2013). As hybridization has traditionally been thought of as a homogenizing force, a major question in evolutionary biology is how speciation can proceed in the face of gene flow (Felsenstein 1981; Bolnick and Fitzpatrick 2007; Feder et al. 2012; Harrison and Larson 2014). Despite a contentious history, it is now recognized that hybridization can actually promote speciation though reinforcement, the process by which enhanced prezygotic isolation is favored in sympatry in response to postzygotic isolation (Dobzhansky 1937; Servedio and Noor 2003; Coyne and Orr 2004). Reinforcement causes reproductive character displacement (RCD), whereby behavioral isolation between two species is heightened in sympatry compared to allopatry. Although multiple different evolutionary forces can lead to such a pattern (reviewed in Hoskin and Higgie 2010), it is considered reinforcement when the mechanism underlying RCD is selection against hybridization (Pfennig and Pfennig 2012). Empirical and theoretical research has indicated that reinforcement may be more common than previously thought (Yukilevich 2012; Hudson and Price 2014), and can both directly finalize speciation in sympatry and indirectly initiate speciation in allopatry (via cascade reinforcement; Ortiz-Barrientos et al. 2009).

Our goal here was to use genomic data to investigate a putative hybrid zone between two species of darters and to examine the strength of multiple postzygotic barriers between these species. The two focal species exhibit a pattern of behavioral isolation consistent with reinforcement of male mating preferences (i.e., male preference for conspecific females is high in allopatry). Whether or not postzygotic isolation is present is unknown. Previous studies have shown a lack of postzygotic isolation through the F1 larval stage. However, the total strength of postzygotic isolation is frequently underestimated by using F1 hybrid inviability as the sole measurement of postzygotic isolation (Wiley et al. 2009; Lemmon and Lemmon 2010). This is particularly problematic because genetic incompatibilities can be masked in F1s due to effects of dominance (Coyne and Orr 2004; Mallet 2006), and maternal provisioning can reduce F1 inviability (Schrader and Travis 2008). Accurate estimates of postzygotic isolation therefore require quantifying postzygotic barriers in F1 adults and in later generation hybrids, but this can be quite challenging in long lived and/or non-model organisms. Measuring the total strength of postzygotic isolation typically necessitates generating multiple generations of hybrid crosses and raising the offspring in the laboratory through the adult life stage. This can be logistically challenging. The current study solves this problem by identifying F1 hybrids in nature and using them to generate second generation hybrids and measure postzygotic isolation.

Darters are a diverse group of stream fishes that have been characterized as a model system for the evolution of speciation via sexual selection. Behavioral isolation evolves before F1 larval inviability in darters (Mendelson 2003; Mendelson et al. 2006, 2007; Williams and Mendelson 2014; Martin and Mendelson 2016a), and there are no known cases of complete F1 inviability through the fertilization and larval hatching stage, even between very distantly related species. The apparent rapid evolution of prezygotic isolation relative to postzygotic isolation in these fish has been attributed to female mate choice on species-specific male color traits (Williams and Mendelson 2010, 2011, 2013). However, recent research in a number of darter species has found that strong conspecific mate preferences are exhibited by males but such preferences are weak (or sometimes absent) in females, and that male coloration functions primarily in male-male competition rather than female mate choice (Zhou et al. 2015; Martin and Mendelson 2016a; Zhou and Fuller 2016; Moran et al. 2017; Mendelson et al. 2018; Moran and Fuller 2018). Thus, males may actually play a stronger role than females in maintaining species boundaries, despite the presence of traditional sex roles and extreme sexual dimorphism.

The present study focuses on the rainbow darter *Etheostoma caeruleum* and the orangethroat darter *Etheostoma spectabile*. The orangethroat darter is a member of the *Ceasia* clade (also referred to as the orangethroat darter clade), which consists of 15 allopatrically distributed species. Time-calibrated gene phylogenies estimate that species within the orangethroat clade last shared a common ancestor 6-7 million years ago (mya) (Bossu et al. 2013). The orangethroat darter clade and rainbow darters are classified together in the subgenus *Oligocephalus.* Divergence time between rainbow and orangethroat darters has been estimated at 22 mya (Near et al. 2011), but these species have very similar male color patterns, ecology, and mating behavior. Thirteen of the orangethroat clade species occur sympatrically with rainbow darters, and ancient hybridization events are evident from the presence of introgressed rainbow darter mitochondrial haplotypes in four orangethroat species (i.e., orangethroat darter *E. spectabile,* current darter *E. uniporum*, brooks darter *E. burri*, and buffalo darter *E. bison*; Ray et al. 2008; Bossu and Near 2009). Molecular evidence also suggests that hybridization is ongoing between the rainbow darter and two species in the orangethroat darter clade (i.e., the buffalo darter and the current darter), as early-generation hybrids have been documented in nature (Bossu and Near 2013; Moran et al. 2017). However, the evolutionary consequences of hybridization in darters remains unexplored.

Recent studies have suggested that selection against interspecific interactions (i.e., mating and fighting) contribute to behavioral isolation between orangethroat and rainbow darters. In sympatric pairings between rainbow darters and five different orangethroat darter clade species, males have been shown to exert strong preferences for mating with conspecific females and fighting with conspecific males (Moran et al. 2017). Such preferences are absent in allopatric pairings of rainbow and orangethroat darters with similar divergence times to the sympatric pairings (Moran and Fuller 2018). This pattern is consistent with both RCD in male mating preferences and divergent agonistic character displacement (ACD) in male fighting preferences. Divergent ACD occurs when selection against interspecific aggressive interactions leads to the evolution of enhanced bias against fighting with heterospecifics in sympatry (Grether et al. 2009). Additionally, behavioral experiments simulating secondary contact between multiple allopatric orangethroat darter clade species revealed that males also prefer to mate and fight with conspecifics over other orangethroat species, but only when they occur sympatrically with rainbow darters (Moran and Fuller 2018). This suggests that RCD and ACD in sympatry between orangethroat and rainbow darters may have cascading effects by incidentally initiating trait evolution and male-driven behavioral isolation among lineages within the orangethroat darter clade. Surprisingly, studies have consistently failed to detect female preferences in orangethroat and rainbow darters for varying components of male color pattern within or between species (Pyron 1995; Fuller 2003; Zhou et al. 2015; Moran et al. 2017).

Whether reinforcement is causing the pattern of RCD in male mating preferences in orangethroat and rainbow darters remains uncertain. Previous investigations into postzygotic barriers between orangethroat and rainbow darters have been limited to examining F1 larval survival, and have found no evidence of hybrid inviability through this life stage (Hubbs and Strawn 1957; Linder 1958; Hubbs 1967; Bossu 2012; Bossu and Near 2013). Here, we use phenotypic and genomic data to confirm that hybridization is ongoing between the orangethroat darter and the rainbow darter, and then investigate postzygotic isolation between these species using both lab-generated and wild-caught hybrids. We test for inviability, sex ratio distortion, sterility, and mating behavioral abnormalities in F1 hybrids, and inviability in backcross hybrids. This represents the most thorough investigation to date into postzygotic isolation in darters. By utilizing natural hybrids, we were able to reveals that postzygotic isolation is much higher than previously thought. We present evidence that hybridization is ongoing and that it is maladaptive, providing critical support for the hypothesis that male-driven behavioral isolation has evolved via reinforcement (and cascade reinforcement) in these species.

## Methods

### Laboratory F1 hybrid cross viability

We first created F1 hybrids in the lab. Adult orangethroat and rainbow darters were collected from two adjacent tributaries of the Vermillion River (Champaign Co., Illinois; Table S1) using a kick seine in April and May 2012. Fish were transported back to the University of Illinois. Crosses were performed by hand-stripping eggs from a single female into a petri dish filled with water from their native stream and subsequently hand-stripping sperm from a single male onto the eggs. Afterward, the water in the petri dish was gently swirled for 1 min to mix the eggs and sperm. Each clutch of eggs was transferred to a separate plastic tub filled with water that was treated with methylene blue (to prevent fungal growth) and stored in an incubator set to 11° C and a 11:14 h light:dark cycle.

Unique male-female pairs were used as parents in each replicate cross. We performed F1 crosses in both direction and “purebred” control crosses with both parental species, with 10–14 replicates per cross type (Table 1). The eggs from each replicate were checked daily for development. As fry hatched, they were transferred to a larger tub in the incubator and fed live brine shrimp nauplii every other day. Fry were transferred out of the incubator and into 19 L and 38 L aquaria at approximately three weeks post hatching. Aquaria were maintained at 19° C and the photoperiod was set to mimic natural daylight hours. After transfer to the aquaria, fish were fed daily *ad libitum* with frozen daphnia and frozen bloodworms.

We measured fertilization success (proportion of eggs that developed pigmented eyes), hatching success (proportion fertilized eggs that yielded free-swimming fry), and larval survival (proportion of hatched eggs that survived to 10 months) of each family. Additionally, to determine whether the mean sex ratio of each cross type deviated from the expected 1:1, we measured the sex ratio of each family after 22 months. By this time, all fish exhibited sexually dimorphic coloration.

All statistical analyses were conducted in R (version 3.4.0). We asked whether each viability metric (fertilization success, hatching success, and larval survival) varied among cross types at the family level using generalized linear models (GLMs), with the viability metric as the independent variable and cross type as the dependent variable. We conducted these analyses using the glm function of the stats package and specified a quasibinomial distribution with logit link function to account for overdispersion in the data. We used the ANOVA function of the car package (Fox 2007) to generate type II analysis of deviance tables and F-tests. We also used one-sample Student’s *t*-tests using the t.test function of the stats package to test whether the proportion of male offspring in a clutch differed from the expected 0.50 in each cross type.

### Backcross viability using wild-caught F1 hybrids

Here, we backcrossed wild-caught F1 hybrid males to orangethroat and rainbow darter females. We used F1 individuals collected from a natural hybrid zone as parents in backcrosses rather than using lab-generated F1s because at two years of age most of our lab-raised orangethroat darters and F1 hybrids failed to engage in mating behavior and females were not gravid. This was not completely unexpected, as orangethroat and rainbow darters can take up to three years to reach sexual maturity in the lab (R. Moran pers. obs.). However, it is also possible that the artificial lab rearing environment lacked a critical cue to trigger the onset of spawning. We therefore only used wild-caught fish for backcrosses.

We collected adult male and female orangethroat and rainbow darters and F1 hybrid males from three tributaries of the Vermillion River (Champaign Co., Illinois; Table S1) in April 2016. We chose to use F1 hybrid males (rather than females) to measure backcross viability because preliminary analyses of our F1 laboratory crosses revealed that: (1) hybrid males are diagnosable due to their color pattern intermediacy between the parental species (see below) (Fig. 1), and (2) the sex ratio of F1 hybrid clutches is dramatically skewed towards males, which suggests that F1 females may be quite rare in natural populations (see below). We confirmed our initial classification of wild-caught fish as orangethroats, rainbows, or hybrids using multivariate phenotypic analyses and genetic sequencing (see below).

**Figure 1.**
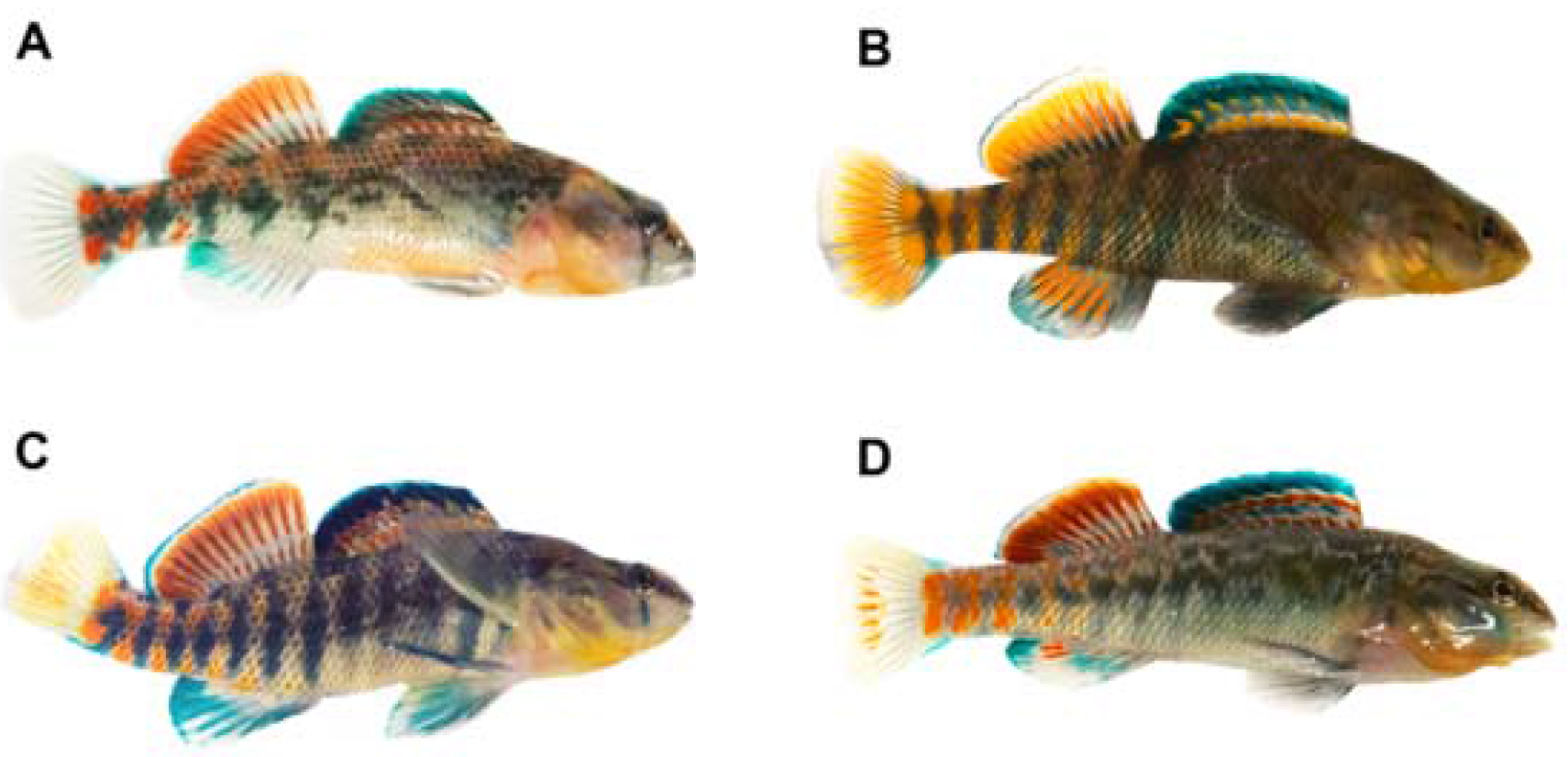
(A) Orangethroat darter and (B) rainbow darter males showing color pattern typical of these species. Orangethroat darters lack the red coloration that is present on the caudal and anal fin in rainbow darters. (C) Wild-caught orangethroat x rainbow darter F1 hybrid male and (D) lab-generated orangethroat x rainbow darter F1 hybrid male showing color pattern characteristics that are combinations of both parental species.

We conducted four cross types with six replicates each. We conducted backcrosses in both directions between the wild-caught F1 males and parental species females and conducted “purebred” control crosses with both parental species (Table 2). Crosses were conducted in breeding aquaria filled with 5-7 cm of naturally colored aquarium gravel, and fluorescent lighting was provided that mimicked the natural photoperiod. Fish were fed frozen bloodworms *ad libitum* each day.

To generate backcrosses, a hybrid male was rotated daily between two 37.9 L breeding aquaria, one of which contained an orangethroat darter female and the other a rainbow darter female. We used a small dip net to rotate hybrid males from one backcross breeding aquarium to the other every day at noon for 14 consecutive days, so that hybrid males spent seven days with each of the two parental females. Eggs were collected from each breeding aquarium immediately after the hybrid male was transferred to the other parental female’s breeding aquarium. Eggs were collected each day for seven days from each purebred parental pair. Purebred parental males were also moved from their breeding aquaria to a separate holding tank for 10 min once a day. During this time, the eggs were collected from the breeding aquaria. Eggs collected from each breeding aquarium were kept together in a 1 L container and maintained as described in the previous section. For each cross, we measured offspring viability at three developmental stages: the proportion of eggs that were fertilized, the proportion of fertilized eggs that survived to hatching, and the proportion of hatched fry that survived to the larval feeding stage (approximately three days post-hatching).

We first asked whether the three measures of viability varied as a function of cross following the same methodology as described above for the F1 crosses. The glht function of the multcomp R package (Hothorn et al. 2017) was used to make post-hoc pairwise comparisons between cross types. We also asked whether females used in backcrosses were as likely to produce eggs as those used in the purebred parental crosses. We conducted two separate Mann-Whitney U tests to determine whether the total number of eggs produced by orangethroat and rainbow darter females differed depending on the identity of the male that they were paired with (i.e., hybrid male or purebred conspecific male). Female standard length did not differ between species (mean ± SE: orangethroat = 61.45 ± 1.43 mm, n = 12; rainbow = 57.97 ± 1.66 mm, n = 12; two-sample *t*-test: t_21.52_= 1.52, p = 0.14), and male standard length did not differ among groups (mean ± SE: hybrids = 65.4 ± 3.2 mm, n = 6; orangethroat = 68.6 ± 3.0 mm, n = 6; rainbow = 68.0 ± 1.6 mm, n = 6; ANOVA: F_2,15_ = 0.34, p = 0.72).

### Wild-caught F1 hybrid male mating and competitive behavior

Both orangethroat and rainbow darters congregate in shallow, gravel riffles of headwater streams during the spring breeding season. Males attempt to guard females by chasing off male competitors and flaring their fins in threat displays. Once a female is ready to spawn, she will perform a nosedig into the gravel and bury herself in the substrate. If multiple males are near the female at this time, male fighting will escalate. One to several males will then attempt to spawn with the female (Winn 1958; Fuller 2003).

We conducted two types of behavioral trials to examine mating behavior of wild-caught F1 hybrid males: dichotomous male choice trials and male competition trials. These behavioral trials used the same wild-caught F1 hybrid males as the backcross experiment described above but used different orangethroat and rainbow darter individuals from the same drainage (i.e., Vermillion River, Champaign Co., Illinois). Previous behavioral studies have shown that orangethroat and rainbow darter males from this drainage exhibit strong preferences for mating and fighting with members of their own species over the other (Zhou and Fuller 2014; Moran et al. 2017; Moran and Fuller 2018). Here, our goal was to ask whether hybrid males show any preference for mating or fighting with members of either parental species. Each behavioral trial involved three fish in a 37.9 L test aquarium positioned under a fluorescent light and filled with 5-7 cm of naturally colored gravel.

For the dichotomous male mate choice trials, a hybrid male was joined by a female orangethroat and a female rainbow darter (n=6). This allowed us to observe whether hybrid males would choose to pursue either female, and if so, whether they exhibited a preference for females of either species. We split each trial into 60 30-s blocks. We scored the number of 30-s blocks in which the male was within one body length of each female for a minimum consecutive time of 5-s (Zhou et al. 2015; Moran and Fuller 2018). We used one-sample Student’s *t*-tests with the t.test function of the stats package in R to test whether the proportion of blocks that the male spent pursuing the orangethroat darter female (versus the total number of blocks spent pursuing either female) differed from the expected 0.50 in each trial.

For the male competition trails, a hybrid male was joined by a male-female pair that were either both orangethroat or both rainbow darters. The goal of these trials was to measure male-male aggressive behavior, but a female was included to elicit male competitive behavior. Each hybrid male participated in two consecutive competition trials, one in which he was joined by an orangethroat darter pair (n=6) and one in which he was joined by a rainbow darter pair (n=6). Thus, each hybrid male was involved in a total of three behavioral trials: one dichotomous male choice trial and two male competition trials. Hybrid males experienced these trial types in random order. Unique purebred fish were used in each trial. We measured hybrid male aggressive behavior by counting the number of attacks (chasing and biting) and fin flares (male threat displays) that the hybrid male performed towards the purebred male in each trial (Zhou et al. 2015; Moran et al. 2017). We asked whether the number of attacks and fin flares that hybrid males directed towards males of the two purebred species differed. We performed GLMs with a negative binomial distribution and logit link function using the glm.nb function of the MASS package in R (Ripley et al. 2017). We performed separate GLM analyses that included the number of male aggressive behaviors (fin flares or attacks) performed in each trial as the dependent variable, and the identity of the purebred species pair in the trial (orangethroat or rainbow) as the independent variable.

### Morphological and histological analyses of testes

To further investigate potential F1 hybrid male sterility, we examined the testes of the six hybrid males and the 12 parental males (six orangethroat and six rainbow darters) that were used in the backcross experiment. Males were euthanized with an overdose of buffered MS-222. We performed gross and histological analyses to compare the testes of the hybrid and purebred males. Testes from each male were fixed in 10% buffered formalin, embedded in paraffin wax, and sectioned. Four μm sections were stained with hematoxylin and eosin and were visually inspected for signs of normal spermatogenesis.

### Color analyses

We used digital photographs to perform multivariate phenotypic analyses of wild-caught orangethroat and rainbow darter males, wild-caught putative F1 hybrid males, and laboratory-generated F1 hybrid males. Our aim was to quantify differences in male color pattern in purebred males and hybrid males, and to statistically verify that hybrid color pattern is distinct and intermediate between purebred species. Such a finding would support our classification of wild-caught F1 hybrid males used in backcross experiments.

We chose to focus on components of male color pattern that differ between the parental species. Superficially, the red and blue banding pattern of orangethroat and rainbow darters looks quite similar, but these species differ in several key ways. Figure 1A and 1B illustrate the differences in male color pattern characteristics between orangethroat and rainbow darters, the most obvious of which are lateral side banding pattern and coloration, anal fin coloration, and caudal fin coloration. Our observations of laboratory-generated and wild-caught F1 hybrid males indicate that hybrids appear to exhibit combinations of both purebred species’ color patterns (Fig. 1C,D).

We measured 36 male color pattern variables (i.e., 27 RGB variable and 9 color proportion variables; see Supplementary Materials for additional details) in the wild-caught hybrid males, orangethroat males, and rainbow males (n=6 each) used in backcross experiments, and in 6 lab-generated F1 hybrid males (which each came from unique families; 3 from ♀ rainbow x ♂ orangethroat crosses, 3 from ♀ orangethroat x ♂ rainbow crosses). We performed Linear Discriminate Analysis (LDA) on the male color pattern data with group (i.e., orangethroat, rainbow, wild-caught hybrid, or laboratory-generated hybrid) as the predictor variable using the lda function of the MASS package in R (Ripley et al. 2017). LDA identifies combinations of independent variables that maximize separation between dependent variables (Mika et al. 1999). Thus, groups with more disparate loadings for a given Linear Discriminant (LD) can be inferred to be more distinct from one another in multivariate signal space. To ask whether male color pattern differs significantly between groups, we conducted multivariate analysis of variance (MANOVA) using the manova function of the stats package in R. Color measurements served as the independent variables and group served as the dependent variable.

### Genotyping wild-caught purebred and hybrid fish

To further verify the purebred or hybrid classification of all fish used in the backcross experiment (42 fish total), we performed single-digest Restriction-site Associated DNA sequencing (RADseq). DNA was isolated from skin and muscle tissue using a modified Puregene protocol. Samples were normalized to a concertation of 15 ng/uL in 50 uL 1x TE. RADseq library preparation with the restriction enzyme *Sbf*I was performed by Floragenex (Eugene, OR, USA), following the methods of Baird et al. (2008). The resulting RADseq library was sequenced as single-end 100 bp reads on two lanes on an Illumina HiSeq 4000 machine.

Sequencing resulted in a total of 37,007,596 reads across the 42 individuals, with a mean ± SE of 881,133 ± 197,553 reads per individual. We used the Stacks (v2.0Beta9; Catchen et al. 2011, 2013) *process_radtags* program to demultiplex samples, remove barcodes, and remove reads of low quality or with ambiguous barcodes. This resulted in a total of 36,232,000 retained reads, which were then supplied to the *denovo_map* pipeline in Stacks to construct a catalog of loci and call SNPs. A minimum of three identical reads were required for each locus (-m 3), with a maximum of three mismatches between loci in each individual (-M 3), and a maximum of two mismatches between loci to be added to the catalog (-n 2). This resulted in a catalog of 63,891 variant sites across 123,901 loci, representing a total of 11,308,200 sites across the genome. The mean ± SE depth of coverage was 23 ± 3X per individual.

The *populations* program in Stacks was used to generate population genetic statistics and to filter loci for analysis of genetic ancestry in Structure (Pritchard et al. 2000). We used *populations* to select loci that were present in all three groups (i.e., orangethroats, rainbows, and putative hybrids) (-p 3) and in at least 50% of the individuals within a group (-r 0.5), with a minimum minor allele frequency of 3%. This filtering resulted in 1,897 SNPs across 1,351 loci (representing a total of 123,472 sites across the genome) for the set of 42 total individuals (6 hybrids, 18 rainbows, 18 orangethroats). To make comparisons between hybrids and parental species, we used *populations* to calculate statistics of genetic differentiation between groups, including SNP-based AMOVA F_ST_ (Weir 1996) and haplotype-based Φ_ST_ (analogous to F_ST_; Excoffier et al. 1992) and D_EST_ (Jost 2008). Unlike F_ST_ and Φ_ST_, D_EST_ is not sensitive to the level of heterozygosity within groups. To obtain an absolute measure of pairwise divergence, we used DnaSP (v6.10.03) (Rozas et al. 2003) to calculated the average number of nucleotide differences between groups (Dxy). To measure the level genetic diversity within groups, we obtained estimates of nucleotide diversity (π), heterozygosity, the percent of polymorphic sites, and the number of private alleles from *populations*.

In the event that more than one SNP was present at a given RAD locus, we only used first SNP for Structure analyses by supplying the --write_single_snp flag to *populations*. This resulted in 1,073 unlinked SNPs that were output in Structure file format. To infer the number of distinct genetic clusters present in the data, we ran Structure with the ancestry model that allowed for admixture and a burnin length of 50,000 followed by 150,000 MCMC repetitions. We performed 20 runs for values of K (i.e., genetic clusters) from 1 to 5 and inferred the optimal value of K using the Evanno method (Evanno et al. 2005) in Structure Harvester (Earl and vonHoldt 2012). Preliminary analyses confirmed the presence of two distinct genetic clusters in the data set, one corresponding to orangethroat darters and the other to rainbow darters (see Results).

To infer the proportion of ancestry associated with orangethroat versus rainbow darters in each hybrid male, we also calculated the hybrid index in GenoDive (v2.0b27) (Meirmans and Van Tienderen 2004) following the method of Buerkle (2005). The hybrid index is a maximum-likelihood estimate of the proportion of alleles in a hybrid individual that originated from one parental species versus the other. We imported the Structure file containing genotype data for 1,073 SNPs across all 42 individuals into GenoDive. For a given hybrid individual, a hybrid index closer to 1 would indicate allele frequencies more similar to that of orangethroat darters, a hybrid index closer to 0 would indicate allele frequencies more similar to that of rainbow darters.

## Results

### Laboratory F1 hybrid cross viability

Fertilization success, hatching success, and larval survival did not differ between F1 hybrid clutches and the “purebred” parental species clutches (Fertilization Success: F_3,39_ = 0.51, p = 0.68; Hatching Success: F_3,39_ = 0.04, p = 0.99; Fry Survival: F_3,32_ = 0.31, p = 0.82; Table 1, Fig. S1). Fertilization success varied greatly across replicate clutches but averaged less than 50% for all cross types. There were five clutches in which none of the eggs developed, possibly due to them being unripe or overly ripe (Moran et al. 2018). Excluding these five crosses from the analysis did not qualitatively change the results. On average, over 50% of fertilized eggs hatched. Mortality was minimal between 10 and 22 months. Eight hybrids and three purebred fish died during this period, but most could be traced to artificial causes (e.g., tank filter failure).

**Table 1.**
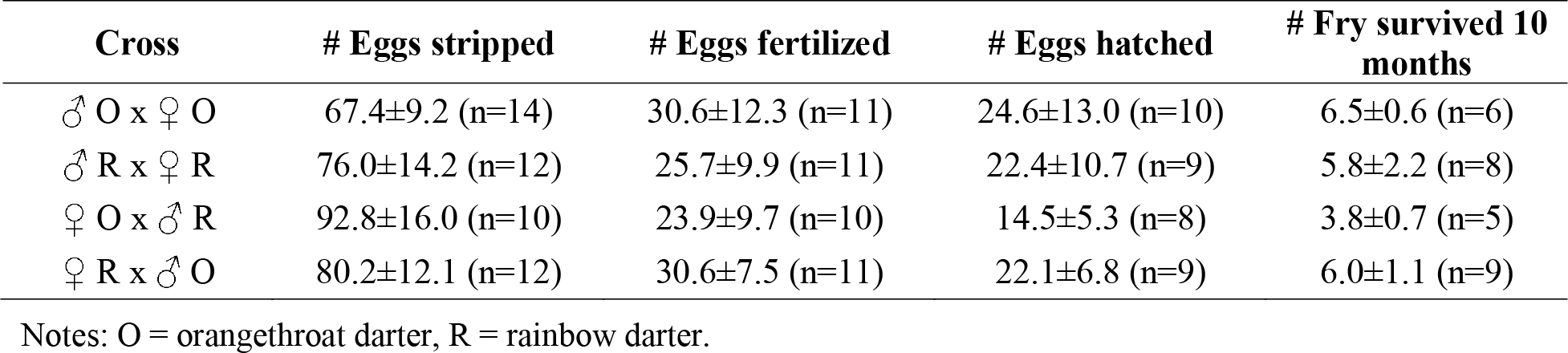
Mean (± standard error) number of total eggs stripped, eggs fertilized, eggs hatched, and fry that survived to 10 months of age in the purebred and hybrid crosses.

| Cross | # Eggs stripped | # Eggs fertilized | # Eggs hatched | # Fry survived 10 months |
| --- | --- | --- | --- | --- |
| ♂ O x ♀ O | 67.4±9.2 (n=14) | 30.6±12.3 (n=11) | 24.6±13.0 (n=10) | 6.5±0.6 (n=6) |
| ♂ R x ♀ R | 76.0±14.2 (n=12) | 25.7±9.9 (n=11) | 22.4±10.7 (n=9) | 5.8±2.2 (n=8) |
| ♀ O x ♂ R | 92.8±16.0 (n=10) | 23.9±9.7 (n=10) | 14.5±5.3 (n=8) | 3.8±0.7 (n=5) |
| ♀ R x ♂ O | 80.2±12.1 (n=12) | 30.6±7.5 (n=11) | 22.1±6.8 (n=9) | 6.0±1.1 (n=9) |

Notes: O = orangethroat darter, R = rainbow darter.

In both F1 hybrid crosses, the sex ratio of the offspring was significantly skewed towards males (♀ orangethroat x ♂ rainbow: mean ± SE proportion male = 0.844 ± 0.104, *t*_5_ = 3.30, p = 0.02; ♀ rainbow x ♂ orangethroat: mean ± SE = 0.948 ± 0.037, *t*_6_ = 12.26, p < 0.00001) (Fig. 2). Only 4 of the 13 F1 hybrid families included females at 22 months. A total of 6 out of 65 F1 hybrids were female. The sex ratio did not differ from the expected 1:1 frequency in purebred crosses (♀ orangethroat x ♂ orangethroat: mean ± SE = 0.594 ± 0.045, *t*_5_ = 2.13, p = 0.09; ♀ rainbow x ♂ rainbow: mean ± SE = 0.450 ± 0.121, *t*_8_ = −0.41, p = 0.69) (Fig. 2). Eleven out of 15 purebred families contained offspring of both sexes at 22 months of age. The total number of offspring per clutch at 22 months did not differ between hybrid and purebred crosses (F_1,26_ = 0.18, p = 0.68).

**Figure 2.**
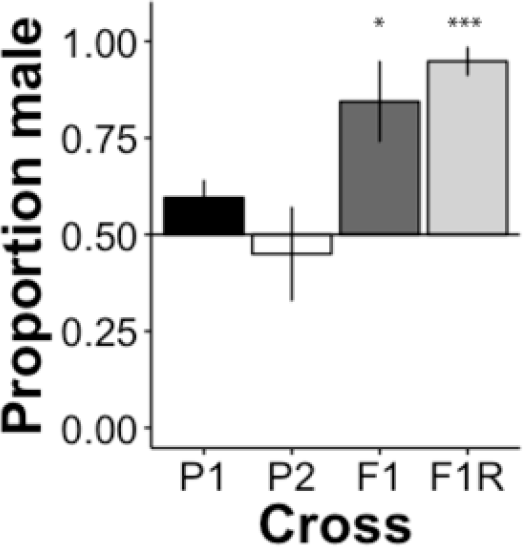
Mean proportion (± standard error) of male offspring in the parental crosses and F1 hybrid crosses at 10 months of age. The deviation from a mean of 0.50 male offspring (i.e., a 1:1 male:female sex ratio) is depicted for each cross type (* = p <0.05, *** = p < 0.001). P1 = ♀ orangethroat x ♂ orangethroat (n=6), P2 = ♀ rainbow x ♂ rainbow (n=8), F1 = ♀ orangethroat x ♂ rainbow (n=5), F1R = ♀ rainbow x ♂ orangethroat (n=9).

### Backcross viability using wild-caught F1 hybrids

Backcrosses suffered higher levels of inviability compared to “purebred” orangethroat and rainbow darter crosses across all three measures of offspring viability (proportion of eggs collected that were fertilized: F_3,20_ = 19.02, p < 0.00001; proportion of fertilized eggs that hatched: F_3,20_ = 3.47, p < 0.05; proportion of hatched eggs that survived to the feeding larval stage: F_3,20_ = 6.95, p < 0.01; Table 2, Fig. 3). Fertilized eggs were collected in 10 out of 12 (83%) of the hybrid male crosses; one hybrid male x orangethroat darter female backcross replicate and one hybrid male x rainbow darter female backcross replicate yielded no fertilized eggs. All purebred crosses produced fertilized eggs. Cumulative survival across all developmental stages was 10X higher in purebred crosses than back-crosses (Table 2). We did not observe any asymmetry in backcross viability: backcross clutches produced by crossing a hybrid male to an orangethroat or rainbow darter female showed equally low levels of viability at each of the three developmental stages measured (Fig. 3). Purebred orangethroat and rainbow darter crosses also did not differ from one another in viability at any stage (Fig. 3).

**Table 2.** Mean (± standard error) number of total eggs collected, eggs fertilized, eggs hatched, and fry that survived to the independently feeding stage in purebred crosses and backcrosses (n = 6 each).

| Cross | # Eggs collected | # Eggs fertilized | # Eggs hatched | # Fry survived to feeding |
| --- | --- | --- | --- | --- |
| ♂ O x ♀ O | 92.33 ± 14.83 | 82.00 ± 12.41 | 61.67 ± 9.90 | 56.17 ± 8.18 |
| ♂ R x ♀ R | 35.50 ± 5.23 | 31.17 ± 3.61 | 21.67 ± 3.19 | 21.00 ± 3.36 |
| ♂ H x ♀ O | 88.00 ± 30.46 | 19.33 ± 14.81 | 10.33 ± 10.14 | 6.00 ± 5.80 |
| ♂ H x ♀ R | 23.17 ± 5.76 | 8.33 ± 2.39 | 2.50 ± 1.77 | 2.17 ± 1.60 |

Notes: O = orangethroat darter, R = rainbow darter, H = F1 hybrid.

**Figure 3.**
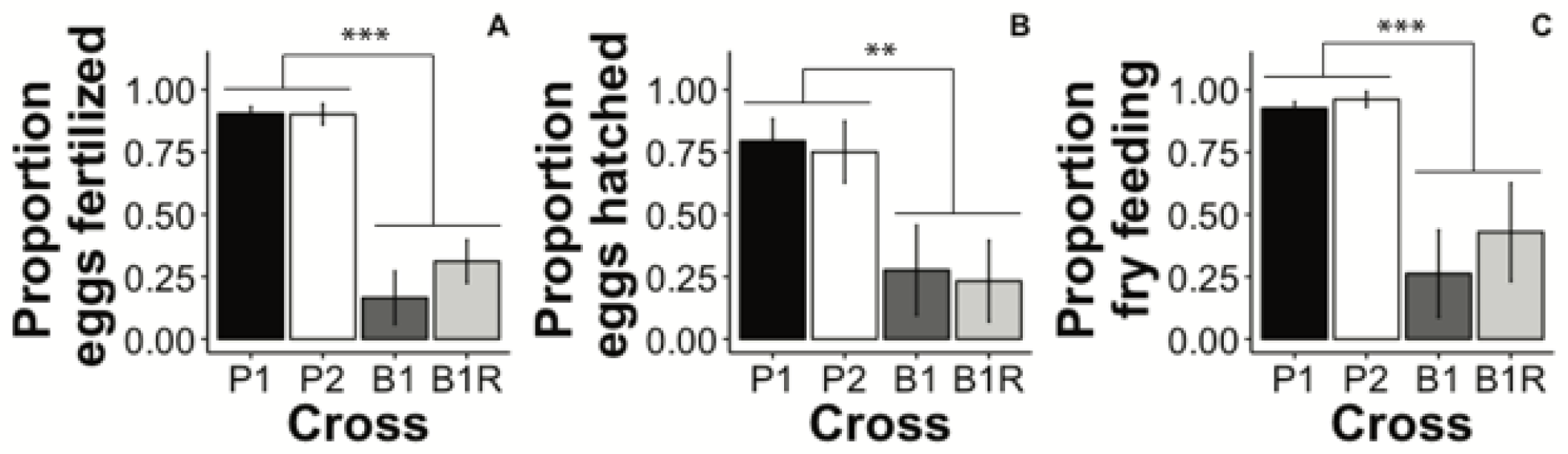
Mean proportion (± standard error) of (A) eggs collected that were fertilized, (B) fertilized eggs that hatched, and (C) hatched fry that survived to the independently feeding stage (approximately three days post-hatching) in the parental crosses and backcrosses (n=6 each). Significance levels are indicated for post-hoc comparisons of purebred crosses and backcrosses (** = p < 0.01, *** = p < 0.001). P1 = ♀ orangethroat x ♂ orangethroat, P2 = ♀ rainbow x ♂ rainbow, B1 = ♀ orangethroat x ♂ F1 hybrid, B1R = ♀ rainbow x ♂ F1 hybrid.

Orangethroat and rainbow darter females used in the crosses produced a similar number of eggs regardless of whether they were paired with a hybrid or a purebred conspecific male (Table 2; orangethroat backcross versus purebred cross: Mann-Whitney U test: U = 18, n = 12, p = 1.00; rainbow backcross versus purebred cross: Mann-Whitney U test: U = 8, n = 12, p = 0.13). In general, rainbow darter females laid fewer, larger eggs compared to orangethroat females, which laid a larger number of smaller eggs (R. Moran pers. obs.). We observed that female orangethroat darters laid two to three times more eggs than female rainbow darters of equivalent size during the duration of this experiment. However, the proportion of offspring surviving through each developmental stage did not differ between species (Fig. 3; Table 2).

### Wild-caught F1 hybrid male mating and competitive behavior

Previous behavioral studies in orangethroat and rainbow darters have shown that males of both species exhibit strong preferences for pursuing females of their own species and preferentially direct aggressive behaviors towards males of their own species (Moran et al. 2017; Moran and Fuller 2018). In contrast, we observed no indication of assortative mating preferences in the wild-caught F1 hybrid males in our dichotomous male choice trials. Hybrid males did not preferentially pursue one purebred species of female over the other (Fig. S2A; *t*_5_ = −0.12, n = 6, p = 0.91). Similarly, hybrid males did not preferentially bias their aggression towards orangethroat or rainbow darter males in the male competition trials. Hybrid males performed a similar number of fin flares (X^2^ = 0.51, n = 6, p = 0.48; Fig. S2B) and attacks (X^2^ = 0.13, n = 6, p = 0.72; Fig. S2C) towards males of both parental species. Additionally, all orangethroat and rainbow darter males engaged in aggressive interactions with the hybrid males.

### Morphological and histological analyses of testes

Gross examination determined that all hybrid males possessed normally developed testes, compared to the purebred orangethroat and rainbow darter males. Comparative histological analysis of the hybrid and purebred male testes revealed that the testes of all males examined contained mature spermatids, and no obvious irregularities in spermatogenesis were observed. Figure S3 shows representative images of testes histology for an orangethroat darter male, a rainbow darter male, and two wild-caught F1 hybrid males.

### Color analyses

The LDA of male color pattern for orangethroat, rainbow, and F1 hybrid males simplified the multivariate color data set of 27 RGB variables and 9 color proportion variables into three LDs. The first two LDs explained a combined total of nearly 87% of the variance in coloration between groups. We visualized the differences in male color pattern among groups in two-dimensional signal space by plotting scores for LD 1 versus LD 2 for each individual (Fig. 4). Orangethroat, rainbow, and F1 hybrid individuals formed tight and well-separated clusters. There was almost complete overlap between the clusters containing the lab-raised and wild-caught F1 hybrid males. Furthermore, hybrid individuals occupied a signal space intermediate between both purebred species along the axis corresponding to LD 1 (Figs. 4, S4).

**Figure 4.**
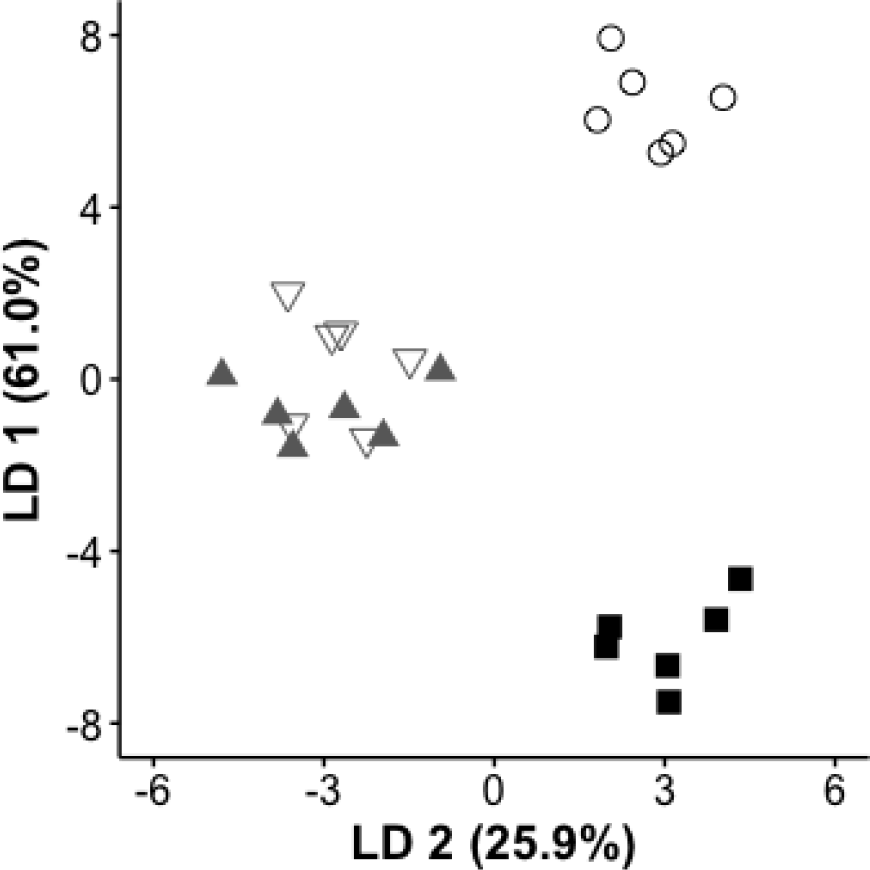
Biplot of the first two Linear Discriminant (LD) axes from the male color pattern LDA of wild-caught orangethroat (▪), wild-caught rainbow (○), wild-caught F1 hybrid males 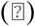, and lab-generated F1 hybird males (▴).

The color proportion measurements had larger LD coefficients compared to the RGB measurements across all three LDs, indicating that the proportion of red and blue coloration on the body and fins are better predictors of group membership than RGB values (Table S2). We therefore used the color proportion measurements for subsequent analyses. There was a significant difference in male color pattern between orangethroat, rainbow, and hybrid males (MANOVA: Pillai’s Trace = 2.33, F_3,20_ = 5.40, p < 0.000001). Male color pattern did not differ between the lab-generated and wild-caught F1 hybrid males (MANOVA: Pillai’s Trace = 0.95, F_1,10_ = 4.22, p = 0.21).

### Genotyping wild-caught purebred and hybrid fish

As expected, notably higher levels of genetic diversity were observed within the hybrid group compared to either parental species (Table 3). Nucleotide diversity (π) and heterozygosity were generally low in both parental species, but higher in rainbow darters compared to orangethroat darters. In the hybrid fish, π was 7.6X higher compared to orangethroat darters and 6.1X higher compared to rainbow darters. Similarly, heterozygosity was 9.8X higher in hybrids compared to orangethroat darters and 5.9X higher in hybrids compared to rainbow darters. The number of private alleles were also an order of magnitude lower in the hybrid group compared to either parental species (Table 3), which is to be expected in F1 hybrids that share half of their alleles with each parental species.

**Table 3.**
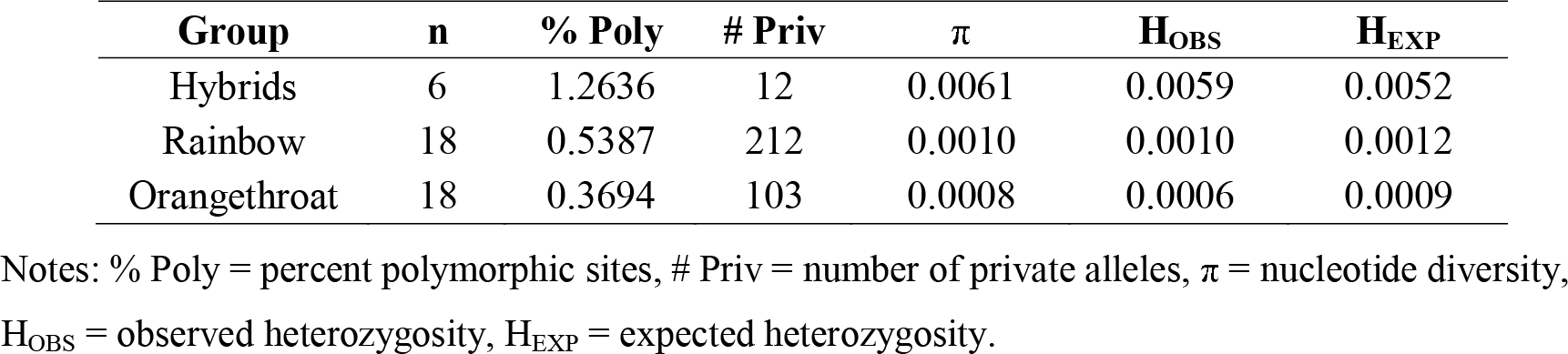
Measurements of genetic diversity within groups (i.e., hybrids, rainbow darters, and orangethroat darters) for all 123,472 sites across 1,351 loci.

| **Group** | **n** | **% Poly** | **# Priv** | **$\pi$** | **H<sub>OBS</sub>** | **H<sub>EXP</sub>** |
| --- | --- | --- | --- | --- | --- | --- |
| Hybrids | 6 | 1.2636 | 12 | 0.0061 | 0.0059 | 0.0052 |
| Rainbow | 18 | 0.5387 | 212 | 0.0010 | 0.0010 | 0.0012 |
| Orangethroat | 18 | 0.3694 | 103 | 0.0008 | 0.0006 | 0.0009 |

Notes: % Poly = percent polymorphic sites, # Priv = number of private alleles, $\pi$ = nucleotide diversity, H<sub>OBS</sub> = observed heterozygosity, H<sub>EXP</sub> = expected heterozygosity.

Patterns of genetic differentiation between groups also supported our classification of hybrid individuals. The SNP-based F_ST_ was lower compared to the haplotype-based Φ_ST_ and D_EST_, but all three measurements of genetic differentiation between groups indicated a high degree of differentiation between orangethroat and rainbow darters, with estimates ranging between 0.689-0.808 (Table 4). As expected, comparisons between hybrids and orangethroat darters and between hybrids and rainbow darters revealed lower levels of differentiation. The average number of nucleotide substitutions per site (Dxy) was 0.01 between orangethroat and rainbow darters. Dxy between hybrids and each of the two parental species was 0.005, exactly half of that between the parental species.

**Table 4.** Measurements of genetic differentiation and divergence between groups (i.e., hybrids, rainbow darters, and orangethroat darters). The SNP-based fixation statistic (F_ST_) was calculated using 1,897 variant sites (SNPs). The haplotype-based fixation statistics (Φ_ST_, D_EST_) and the average number of nucleotide substitutions per site (D_XY_) were calculated using 123,472 sites across 1,351 loci.

| Comparison | $F_{ST}$ | $\Phi_{ST}$ | $D_{EST}$ | $D_{XY}$ |
| --- | --- | --- | --- | --- |
| Hybrids - Rainbows | 0.327 | 0.495 | 0.305 | 0.005 |
| Hybrids - Orangethroats | 0.315 | 0.454 | 0.271 | 0.005 |
| Rainbows - Orangethroats | 0.689 | 0.792 | 0.808 | 0.010 |

The Structure analysis of 1,073 SNPs present in the set of 42 individuals used in the backcross experiment revealed an optimal K of 2 according to the Evanno method implemented in Structure Harvester (Table S3). As with the color analyses, the genetic analyses confirmed our original diagnosis of the wild-caught orangethroat darters, rainbow darters, and F1 hybrid males that were used in the backcross experiment (Fig. 5). With K set to 2, the 18 orangethroat darter individuals were assigned 98% membership to cluster 1, and the 18 rainbow darter individuals were assigned 99% membership to cluster 2. The assignments of the six hybrid males were split between clusters and averaged 53% membership to the orangethroat cluster and 47% membership to the rainbow cluster. The hybrid index scores calculated for the hybrids yielded qualitatively similar results; the maximum likelihood estimate for the proportion of orangethroat darter ancestry in each hybrid male ranged from 0.501-0.566 (Table S4).

**Figure 5.**
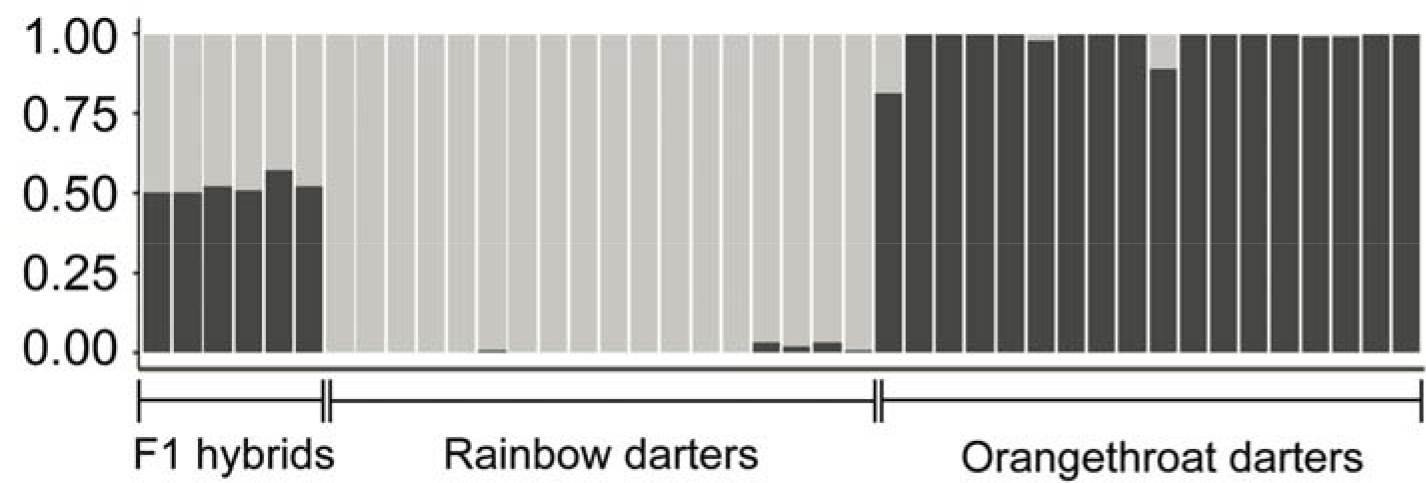
Probability of membership to the rainbow darter cluster (light gray) or orangethroat darter cluster (dark gray) for each individual used in the backcross experiment. Structure analysis was conducted using 1,073 SNPs with the number of clusters (K) set to 2.

## Discussion

Our results have broad implications for addressing the problem of measuring postzygotic isolation in non-model systems and for investigating the selective forces that drive speciation in darters. In many systems, reinforcement has been discounted because behavioral isolation is stronger than F1 inviability or sterility among recently diverged lineages (e.g., birds: Price and Bouvier 2002; Lijtmaer et al. 2003; fish: Mendelson 2003; Van Der Sluijs et al. 2008; frogs: Sasa et al. 2008; Malone and Fontenot 2008; insects: Rull et al. 2013; Gray et al. 2016). Yet quantitative geneticists have known for decades that second generation hybrids (F2s and backcrosses) often have greatly reduced survival (Stebbins 1958; Lynch 1991). Assessing the fitness of second generation hybrids is difficult in many taxa due to long generation times and the need to develop in a ‘natural’ environment (Wiley et al. 2009; Lemmon and Lemmon 2010). We present a strong strategy for investigating postzygotic isolation in long lived, non-model organisms by using morphological and genomic data to identify F1s in nature and then utilizing them to facilitate measurements of postzygotic isolation that effect the adult life stage and hybrid generations. Importantly, our findings drastically change how we think about speciation in darters. Despite being among the most highly diverse groups of vertebrates in North America, our understanding of how speciation occurs in these fishes is incomplete due to the difficulties of performing breeding studies. Below we discuss the unexpectedly high degree of postzygotic isolation that we observed between orangethroat and rainbow darters and its implications for male-driven speciation via reinforcement and cascade reinforcement.

### Patterns of F1 and backcross hybrid inviability

We found high levels of postzygotic isolation between orangethroat and rainbow darters in the form of multiple isolating barriers spanning across hybrid life stages and generations. This system was previously thought to lack substantial postzygotic isolation among species due to high survival of F1 larvae compared to purebred crosses (Hubbs and Strawn 1957; Hubbs 1959). Our results corroborated these previous findings. Clutches resulting from F1 crosses did not exhibit reduced fertilization, hatchability, or survival through adulthood compared to purebred crosses. However, we did observe dramatically distorted sex ratios in F1 crosses. Heterospecific crosses in both directions were heavily skewed towards males. Clutches from purebred crosses did not deviate from a 1:1 sex ratio, and most natural darter populations have also been shown to maintain 1:1 sex ratios in adults (Page 1983). Whether the male-skewed sex ratio in F1 hybrids creates selection favoring assortative mating and behavioral isolation in areas of sympatry is unclear. Such a scenario may be present in *Neochromis* cichlids, which appear to have evolved assortative mating among incipient species in response to sex ratio distortion in hybrid clutches (Seehausen et al. 1999). The mechanisms underlying the lack of adult F1 females is also unknown. Investigation into the genetics of sex determination in darters would add insight into why female hybrids are missing from F1 hybrid clutches.

We also documented substantial postzygotic isolation between orangethroat and rainbow darters in the backcross generation. When wild-caught F1 males were crossed to females of both parental species, backcross clutches in both directions had dramatically reduced fertilization success, hatching success, and larval survival compared to clutches resulting from purebred parental crosses. The dramatic reduction in fertilization success observed in the backcross clutches is likely attributable to genetic incompatibilities being unmasked in backcross progeny, rather than F1 hybrid male sterility. Wild-caught F1 hybrid males did not exhibit any morphological or histological defects of the testes, and fertilized eggs in 10 out of 12 backcrosses. Many of the backcrosses also produced embryos with obvious developmental abnormalities that died before hatching (R. Moran pers. obs.). A small number of progeny resulting from backcrosses to both parental species were able to survive until the free feeding larval stage, indicating that although intrinsic postzygotic isolation between orangethroat and rainbow darters is very high, it is not complete. This has implications for the evolution of mating preferences in this system, which previous studies have shown to be consistent with reinforcement (see below) (Moran et al. 2017; Moran and Fuller 2018).

### Implications for reinforcement

Genome-wide sequence data indicated high genetic differentiation and a 1% nucleotide divergence between orangethroat and rainbow darters. Heterozygosity and nucleotide diversity were generally low in both species, but higher in rainbow darters. This observation is consistent with previous analyses of genetic diversity in these species (Moran et al. 2017), and may reflect higher levels of population connectivity in rainbow darters compared to orangethroat darters (Page 1983).

Notably, our results indicate that F1 hybrids form in nature, and that we can accurately diagnose hybrid males based on color pattern attributes that are intermediate between the two purebred parental species. Molecular markers have also been used to document the presence of naturally occurring F1s, F2s, and backcrosses in both directions between rainbow darters and the orangethroat darter clade species *E. bison* (the buffalo darter) (Bossu and Near 2013), and F1 hybrids between rainbow darters and the orangethroat darter clade species *E. uniporum* (the current darter) (Moran et al. 2017).

The evidence for contemporaneous hybridization between orangethroat and rainbow darters together with the high levels of postzygotic isolation observed provide critical support for previous claims that reinforcement is responsible for driving the patterns of RCD (and potentially ACD) documented in this system (Zhou and Fuller 2014; Moran et al. 2017; Moran and Fuller 2018). Orangethroat and rainbow darter males from sympatric populations consistently show strong biases for mating with conspecific females and fighting with conspecific males (when given a choice between orangethroat or rainbow darters). Such biases are not present in orangethroat and rainbow darters that occur in allopatry with respect to one another (Moran and Fuller 2018). The presence of strong postzygotic isolation and ongoing hybridization between these species has likely created selection favoring the high levels of behavioral isolation observed in sympatry compared to allopatry. Selection to avoid interspecific male-male aggressive interactions in sympatric populations (i.e., ACD) presumably acts to facilitate the co-occurrence of these species in such close proximity to one another in riffle microhabitats during the spawning season. In turn, the fact that orangethroat and rainbow darters occur syntopically on the same spawning grounds increases the potential for hybridization, which can then further fuel RCD via reinforcement. In this manner RCD and ACD may act in a positive feedback loop to strengthen male behavioral biases against heterospecific females and males (Vallin et al. 2012; Moran and Fuller 2018).

The lack of behavioral biases in wild-caught F1 males stands in contrast to the strong biases that were previously documented for sympatric male orangethroat and rainbow darters from the same drainage (Moran et al. 2017; Moran and Fuller 2018). Wild-caught F1 hybrid males pursued females of both parental species equally and engaged in a comparable amount of aggressive interactions with males of both parental species. Similarly, females and males of both parental species did not show any mating or fighting biases against hybrid males. These observations suggest that F1 males are behaviorally intermediate between the two parental species, similar to the pattern we observed in male color pattern. Furthermore, it has previously been argued that in sympatry, selection favors males who fight with conspecific males (over access to conspecific females) and ignore heterospecific males, in order to avoid costly, unnecessary aggression (Moran et al. 2017; Moran and Fuller 2018). The fact that F1 males engage in contests with males of both parental species suggests that they may pay the costs associated with increased fighting by engaging males of both species.

Evidence from the present study also supports the hypothesis that cascade reinforcement is responsible for the surprisingly high levels of male-driven behavioral isolation present between species within the orangethroat clade (Moran and Fuller 2018). By promoting the evolution of mating traits, reinforcement between two species can incidentally cause behavioral isolation among populations within a single species, termed cascade reinforcement (reviewed in Comeault and Matute 2016). Overtime, cascade reinforcement can cause isolated populations within one species that is experiencing reinforcement with a close relative to diverge to such an extent that they are considered distinct species. We hypothesize that such a phenomenon is occurring in orangethroat darters as a correlated effect of reinforcement with rainbow darters. Males from orangethroat clade species that do not co-occur with one another but do occur sympatrically with rainbow darters exert strong preferences for conspecific over heterospecific orangethroat darter females (Moran et al. 2017). It is possible that the parallel occurrence of reinforcement selecting for increased behavioral isolation between sympatric rainbow darters and multiple species within the orangethroat clade has incidentally led to mismatches in mating preferences and behavioral isolation between species within the orangethroat clade. The alternative hypothesis that sexual selection within species is responsible for this pattern is unlikely, as populations of orangethroat darters that are allopatric from other species in the orangethroat darter clade and from rainbow darters have no detectable levels of behavioral isolation.

### Conclusions

We used genomic data to demonstrate that hybridization is ongoing between orangethroat and rainbow darters. These species were previously thought to lack substantial postzygotic isolation, but we observed dramatically skewed sex ratios in F1s and a high degree of inviability in backcrosses. The results of this study demonstrate that selection to avoid hybridization may be more important than previously thought in darters. Our findings also inform our understanding of how speciation occurs in a highly diverse vertebrate group with traditional sex roles and dimorphism but no apparent female mate preferences. Darters provide a unique example of how male preferences alone can promote mating and fighting trait evolution concurrently between sympatric and allopatric lineages. The extensive amount of postzygotic isolation present between orangethroat and rainbow darters suggests that reinforcement promotes the previously documented patterns of RCD in male mating preferences between these species (Moran and Fuller 2018). Furthermore, this implies that cascade effects of reinforcement may be responsible for the evolution of male-driven behavioral isolation between recently diverged lineages within the orangethroat darter clade that occur sympatrically with rainbow darters (Moran et al. 2017). Darters provide an intriguing study system for future investigations into the genetics/genomics of hybridization, reinforcement, and speciation.

## Acknowledgements

The treatment of animals used in this study was in compliance with the University of Illinois Institutional Animal Care and Use Committee (IACUC) under protocols #12055 and #17031. Collection of wild fish was approved by the Illinois Department of Natural Resources under Scientific Collecting Permits A12.4035 and A16.4035. We thank Adam Stern (Veterinary Diagnostic Laboratory, University of Illinois) for assistance with gamete histology, and Jason Boone (Floragenex) for logistic support with RAD library preparation and sequencing. This research was supported by funding from the National Science Foundation (DEB 0953716, DEB 1210743, DGE 1069157, and IOS 1701676), the United States Department of Agriculture (Cooperative State Research, Education, and Extension Service project number ILLU 875-952), and the University of Illinois at Urbana-Champaign.

## References

Abbott, R., D. Albach, S. Ansell, J. W. Arntzen, S. J. E. Baird, N. Bierne, J. Boughman, A. Brelsford, C. A. Buerkle, R. Buggs, R. K. Butlin, U. Dieckmann, F. Eroukhmanoff, A. Grill, S. H. Cahan, J. S. Hermansen, G. Hewitt, A. G. Hudson, C. Jiggins, J. Jones, B. Keller, T. Marczewski, J. Mallet, P. Martinez-Rodriguez, M. Möst, S. Mullen, R. Nichols, A. W. Nolte, C. Parisod, K. Pfennig, A. M. Rice, M. G. Ritchie, B. Seifert, C. M. Smadja, R. Stelkens, J. M. Szymura, R. Väinölä, J. B. W. Wolf, and D. Zinner. 2013. Hybridization and speciation. J. Evol. Biol. 26:229–246.

Baird, N. A., P. D. Etter, T. S. Atwood, M. C. Currey, A. L. Shiver, Z. A. Lewis, E. U. Selker, W. A. Cresko, and E. A. Johnson. 2008. Rapid SNP Discovery and Genetic Mapping Using Sequenced RAD Markers. PLoS One 3:e3376.

Bolnick, D. I., and B. M. Fitzpatrick. 2007. Sympatric Speciation: Models and Empirical Evidence. Annu. Rev. Ecol. Evol. Syst. 38:459–487.

Bossu, C. M. 2012. Linking population genetic patterns of introgressive hybridization to the evolution of reproductive isolating barriers in darters (Percidae). Yale University.

Bossu, C. M., J. M. Beaulieu, P. A. Ceas, and T. J. Near. 2013. Explicit tests of palaeodrainage connections of southeastern North America and the historical biogeography of Orangethroat Darters (Percidae: Etheostoma: Ceasia). Mol. Ecol. 22:5397–5417.

Bossu, C. M., and T. J. Near. 2013. Characterization of a contemporaneous hybrid zone between two darter species (Etheostoma bison and E. caeruleum) in the Buffalo River System. Genetica 141:75–88.

Bossu, C. M., and T. J. Near. 2009. Gene trees reveal repeated instances of mitochondrial DNA Introgression in orangethroat darters (percidae: etheostoma). Syst. Biol. 58:114–129.

Buerkle, C. A. 2005. Maximum-likelihood estimation of a hybrid index based on molecular markers. Mol. Ecol. Notes 5:684–687.

Catchen, J., P. A. Hohenlohe, S. Bassham, A. Amores, and W. A. Cresko. 2013. Stacks: an analysis tool set for population genomics. Mol. Ecol. 22:3124–3140.

Catchen, J. M., A. Amores, P. Hohenlohe, W. Cresko, and J. H. Postlethwait. 2011. Stacks: building and genotyping Loci de novo from short-read sequences. G3 (Bethesda). 1:171–82.

Coyne, J., and H. Orr. 2004. Speciation. Sunderland, MA.

Dobzhansky, T. 1937. Genetics and the Origin of Species. Columbia university press.

Earl, D. A., and B. M. vonHoldt. 2012. STRUCTURE HARVESTER: a website and program for visualizing STRUCTURE output and implementing the Evanno method. Conserv. Genet. Resour. 4:359–361.

Evanno, G., S. Regnaut, and J. Goudet. 2005. Detecting the number of clusters of individuals using the software structure: a simulation study. Mol. Ecol. 14:2611–2620.

Excoffier, L., P. Smouse, and J. Quattro. 1992. Analysis of molecular variance infered from metric distances among DNA haplotypes: application to human mitochondrial DNA restricyion data. Genetics 131:479–491.

Feder, J. L., S. P. Egan, and P. Nosil. 2012. The genomics of speciation-with-gene-flow. Trends Genet. 28:342–350.

Felsenstein, J. 1981. Skepticism towards santa rosalia, or why are there so few kinds of animals? Evolution. 35:124.

Fox, J. 2007. Package “car”. R Foundation for Statistical Computing.

Fuller, R. C. 2003. Disentangling female mate choice and male competition in the rainbow darter, Etheostoma caeruleum. Copeia 2003:138–148.

Gray, D. A., N. J. Gutierrez, T. L. Chen, C. Gonzalez, D. B. Weissman, and J. A. Cole. 2016. Species divergence in field crickets: Genetics, song, ecomorphology, and pre- and postzygotic isolation. Biol. J. Linn. Soc. 117:192–205.

Grether, G. F., N. Losin, C. N. Anderson, and K. Okamoto. 2009. The role of interspecific interference competition in character displacement and the evolution of competitor recognition. Biol. Rev. 84:617–635.

Harrison, R. G., and E. L. Larson. 2014. Hybridization, Introgression, and the Nature of Species Boundaries. J. Hered. 105:795–809. Oxford University Press.

Heins, D. C., J. A. Baker, and D. J. Tylicki. 1996. Reproductive season, clutch size, and egg size of the rainbow darter, Etheostoma caeruleum, from the Homochitto River, Mississippi, with an evaluation of data from the literature. Copeia 1996:1005–1010.

Hendry, A. P., D. I. Bolnick, D. Berner, and C. L. Peichel. 2009. Along the speciation continuum in sticklebacks. J. Fish Biol. 75:2000–2036.

Hoskin, C. J., and M. Higgie. 2010. Speciation via species interactions: The divergence of mating traits within species. Ecol. Lett. 13:409–420.

Hothorn, T., F. Bretz, P. Westfall, and R. Heiberger. 2017. Package “multcomp.”

Hubbs, C. 1967. Geographic variations in survival of hybrids between etheostomatine fishes. Texas Meml. Museum Bull. 72pp.

Hubbs, C. 1959. Laboratory hybrid combinations among etheostomatine fishes. Tex. J. Sci. 11:49–56.

Hubbs, C., and K. Strawn. 1957. Relative variability of hybrids between the Darters, Etheostoma spectabile and Percina caprodes. Evolution. 11:1–10.

Hudson, E. J., and T. D. Price. 2014. Pervasive reinforcement and the role of sexual selection in biological speciation. J. Hered. 105:821–833.

Jost, L. 2008. GST and its relatives do not measure differentiation. Mol. Ecol. 17:4015–4026.

Kitano, J., S. Mori, and C. L. Peichel. 2007. Phenotypic divergence and reproductive isolation between sympatric forms of Japanese threespine sticklebacks. Biol. J. Linn. Soc. 91:671–685.

Lemmon, E. M., and A. R. Lemmon. 2010. Reinforcement in chorus frogs: Lifetime fitness estimates including intrinsic natural selection and sexual selection against hybrids. Evolution. 64:1748–1761.

Lijtmaer, D. A., B. Mahler, and P. L. Tubaro. 2003. Hybridization and postzygotic isolation patterns in pigeons and doves. Evolution. 57:1411.

Linder, A. D. 1958. Behavior and hybridization of two species of Etheostoma (Percidae). Trans. Kansas Acad. Sci. 61:195–212.

Lynch, M. 1991. The genetic interpretation of inbreeding depression and outbreeding depression. Evolution. 45:622–629.

Mallet, J. 2005. Hybridization as an invasion of the genome. Trends Ecol. Evol. 20:229–237.

Mallet, J. 2006. What does Drosophila genetics tell us about speciation?

Malone, J. H., and B. E. Fontenot. 2008. Patterns of Reproductive Isolation in Toads. PLoS One 3:e3900.

Martin, M. D., and T. C. Mendelson. 2016a. Male behaviour predicts trait divergence and the evolution of reproductive isolation in darters (Percidae: Etheostoma). Anim. Behav. 112:179–186.

Martin, M. D., and T. C. Mendelson. 2016b. The accumulation of reproductive isolation in early stages of divergence supports a role for sexual selection. J. Evol. Biol. 29:676–689.

Meirmans, P., and P. H. Van Tienderen. 2004. GENOTYPE and GENODIVE: two\nprograms for the analysis of genetic diversity of asexual organisms. Mol. Ecol. Notes 792–794.

Mendelson, T. C. 2003. Sexual isolation evolves faster than hybrid inviability in a diverse and sexually dimorphic genus of fish (Percidae: Etheostoma). Evolution. 57:317–327.

Mendelson, T. C., J. M. Gumm, M. D. Martin, and P. J. Ciccotto. 2018. Preference for conspecifics evolves earlier in males than females in a sexually dimorphic radiation of fishes. Evolution. 72:337–347.

Mendelson, T. C., V. E. Imhoff, and M. K. Iovine. 2006. Analysis of early embryogenesis in Rainbow and Banded Darters (Percidae: Etheostoma) reveals asymmetric postmating barrier. Environ. Biol. Fishes 76:351–360.

Mendelson, T. C., V. E. Imhoff, and J. J. Venditti. 2007. The accumulation of reproductive barriers during speciation: Postmating barriers in two behaviorally isolated species of darters (Percidae: Etheostoma). Evolution. 61:2596–2606.

Mika, S., G. Ratsch, J. Weston, B. Scholkopf, and K. R. Mullers. 1999. Fisher discriminant analysis with kernels. Proc. 1999 IEEE Signal Process. Soc. Work. 41–48.

Moran, R. L., and R. C. Fuller. 2018. Male-driven reproductive and agonistic character displacement in darters and its implications for speciation in allopatry. Curr. Zool. 64:101–113.

Moran, R. L., M. Zhou, J. M. Catchen, and R. C. Fuller. 2017. Male and female contributions to behavioral isolation in darters as a function of genetic distance and color distance. Evolution. 71:2428–2444.

Moran, R. L., R. M. Soukup, M. Zhou, and R. C. Fuller. 2018. Egg viability decreases rapidly with time since ovulation in the rainbow darter Etheostoma caeruleum◻: implications for the costs of choosiness. J. Fish Biol. 92:532–536.

Near, T. J., C. M. Bossu, G. S. Bradburd, R. L. Carlson, R. C. Harrington, P. R. Hollingsworth, B. P. Keck, and D. A. Etnier. 2011. Phylogeny and temporal diversification of darters (Percidae: Etheostomatinae). Syst. Biol. 60:565–595.

Ortiz-Barrientos, D., A. Grealy, and P. Nosil. 2009. The genetics and ecology of reinforcement: implications for the evolution of prezygotic isolation in sympatry and beyond. Ann. N. Y. Acad. Sci. 1168:156–82.

Page, L. 1983. Handbook of darters.

Pfennig, D., and K. Pfennig. 2012. Evolution’s wedge: competition and the origins of diversity.

Price, T. D., and M. M. Bouvier. 2002. The evolution of F1 postzygotic incompatibilities in birds. Evolution. 56:2083–2089.

Pritchard, J. K., M. Stephens, and P. Donnelly. 2000. Inference of population structure using multilocus genotype data. Genetics 155:945–59.

Pyron, M. 1995. Mating patterns and a test for female mate choice in Etheostoma spectabile (Pisces, Percidae). Behav. Ecol. Sociobiol. 36:407–412.

Ray, J. M., N. J. Lang, R. M. Wood, and R. L. Mayden. 2008. History repeated: recent and historical mitochondrial introgression between the current darter Etheostoma uniporum and rainbow darter Etheostoma caeruleum (Teleostei: Percidae). J. Fish Biol. 72:418–434.

Ripley, B., B. Venables, D. Bates, K. Hornik, and A. Gebhardt. 2017. Package “MASS.”

Rozas, J., J. C. Sanchez-DelBarrio, X. Messeguer, and R. Rozas. 2003. DnaSP, DNA polymorphism analyses by the coalescent and other methods. Bioinformatics 19:2496–2497.

Rull, J., S. Abraham, A. Kovaleski, D. F. Segura, M. Mendoza, M. C. Liendo, and M. T. Vera. 2013. Evolution of pre-zygotic and post-zygotic barriers to gene flow among three cryptic species within the Anastrepha fraterculus complex. Entomol. Exp. Appl. 148:213–222.

Rundle, H. D., and M. C. Whitlock. 2001. A genetic interpretation of ecologically dependent isolation. Evolution 55:198–201.

Sasa, M. M., P. T. Chippindale, and N. A. Johnson. 2008. Patterns of postzygotic isolation in frogs. Wiley Online Libr. 52:1811–1820.

Schrader, M., and J. Travis. 2008. Testing the Viviparity-Driven-Conflict Hypothesis: Parent-Offspring Conflict and the Evolution of Reproductive Isolation in a Poeciliid Fish. Source Am. Nat. 172:806–817.

Seehausen, van Alphen, and Lande. 1999. Color polymorphism and sex ratio distortion in a cichlid fish as an incipient stage in sympatric speciation by sexual selection. Ecol. Lett. 2:367–378.

Servedio, M. R., and M. A. F. Noor. 2003. The Role of Reinforcement in Speciation: Theory and Data. Annu. Rev. Ecol. Evol. Syst. 34:339–364.

Stebbins, G. L. 1958. The inviability, weakness, and sterility of interspecific hybrids. Adv. Genet. 9:147–215.

Stelkens, R. B., K. A. Young, and O. Seehausen. 2010. The accumulation of reproductive incompatibilities in African cichlid fish. Evolution. 64:617–633.

Vallin, N., A. M. Rice, R. I. Bailey, A. Husby, and A. Qvarnström. 2012. Positive feedback between ecological and reproductive character displacement in a young avian hybrid zone. Evolution. 66:1167–1179.

Van Der Sluijs, I., T. J. M. Van Dooren, O. Seehausen, and J. J. M. Van Alphen. 2008. A test of fitness consequences of hybridization in sibling species of Lake Victoria cichlid fish. J. Evol. Biol. 21:480–91. Blackwell Publishing Ltd.

Weir, B. S. and Cockerham, C. 1996. Genetic data analysis II: Methods for discrete population genetic data. Sinauer Assoc. Inc., Sunderland, MA, USA.

Wiley, C., A. Qvarnström, G. Andersson, T. Borge, and G.-P. Saetre. 2009. Postzygotic Isolation over Multiple Generations of Hybrid Descendents in a Natural Hybrid Zone: How Well Do Single-Generation Estimates Reflect Reproductive Isolation? Source Evol. 63:1731–1739.

Williams, T. H., and T. C. Mendelson. 2010. Behavioral Isolation Based on Visual Signals in a Sympatric Pair of Darter Species. Ethology 116:1038–1049.

Williams, T. H., and T. C. Mendelson. 2011. Female preference for male coloration may explain behavioural isolation in sympatric darters. Anim. Behav. 82:683–689.

Williams, T. H., and T. C. Mendelson. 2013. Male and female responses to species-specific coloration in darters (Percidae: Etheostoma). Anim. Behav. 85:1251–1259.

Williams, T. H., and T. C. Mendelson. 2014. Quantifying Reproductive Barriers in a Sympatric Pair of Darter Species. Evol. Biol. 41:212–220.

Winn, H. E. 1958. Observations on the Reproductive Habits of Darters (Pisces-Percidae). Source Am. Midl. Nat. 59:190–212. The University of Notre Dame.

Yukilevich, R. 2012. Asymmetrical patterns of speciation uniquely support reinforcement in drosophila. Evolution. 66:1430–1446.

Zhou, M., and R. C. Fuller. 2016. Intrasexual competition underlies sexual selection on male breeding coloration in the orangethroat darter, Etheostoma spectabile. Ecol. Evol. 6:3513–3522.

Zhou, M., E. R. Loew, and R. C. Fuller. 2015. Sexually asymmetric colour-based species discrimination in orangethroat darters. Anim. Behav. 106:171–179.

